# Plant Growth analysis using computer. An auxiliary computational program. II

**DOI:** 10.1101/2024.04.02.587807

**Authors:** Tomás de Aquino Portes

## Abstract

The result of photosynthetic activity by plants is mass gain. Essential mineral elements are immobilized in the plant mass, the main element being carbon. In photosynthesis, this element is linked to one another forming carbon chains and to link one carbon to another the plant uses solar energy, making the mass rich in energy. Therefore, quantifying plant mass is interesting to estimate the efficiency of plants in accumulating mass and energy. Mass quantification is done using the growth analysis technique. The objective of this work was to develop a computer program to assist in the plant growth analysis. It consisted of adjusting mathematical equations to data on leaf area, total and leaf dry mass. Once the best equations were found, the instantaneous growth indicators, CGR, RGR, NAR, SLA, LAR, LAD and carbon flux (pn) were estimated from them. The adjusted equations were linear, logistic, quadratic, cubic, quadratic exponential and cubic exponential. The equations that best fit to the data were the quadratic exponential and cubic exponential. The program was developed in the C* language. For comparison purposes, the average physiological indicators of growth were estimated using the classic equations found in the literature.

**Highlight:** Plant growth analysis is a very useful technique for evaluating plant mass gain. To facilitate the analysis, a computer program presented in this article was developed. The software is important because there is nothing similar available and will facilitate the work of researchers working on the subject

## Introduction

The difference between CO_2_ fixed photosynthetically and that lost in respiration and photorespiration represents a net gain in mass by the plant resulting in growth. This mass is distributed among the various plant organs such as leaves, stems, branches, roots, inflorescences and fruits used in human and animal nutrition and also produces energy and enriches and protects the soil with organic matter. The difference, when positive, represents carbon sequestration, contributing to the reduction of its excess in the atmosphere, mitigating the impact of the very worrying global warming (Cox *et al*., 2000; IPCC, 2014; IPCC, 2013). Therefore, quantifying this mass gain is of great economic and ecological interest. Quantifying the difference between the amount of CO_2_ fixed by photosynthesis and that lost through respiration and photorespiration is done using the growth analysis technique (Kvet *et al*., 1971; Portes *et al*., 2022).

Growth can be analyzed from several aspects. For example: measuring the growth of the plant in height, or the growth of its diameter, or even of parts or organs of the plant (allometric growth). The most common, however, is the measurement of the phytomass growth of plant communities, in greenhouses or, in field conditions.

Although it is a classic technique (Blackman, 1919), plant growth analysis is still, and will continue to be, of great use in quantifying growth and determining the partitions of photosynthates between the various plant organs, providing support for those working towards grain or forage production, helping to select the best genotypes for such purposes, quantify carbon sequestration, and plant responses to the application of a product to the soil or to the plant itself. It is a technique that requires few instrumental resources and is therefore low-cost, making it an excellent tool for researchers in developing countries (Kvet *et al*., 1971; Portes and Carvalho, 2009; Portes *et al*., 2022; Westhoff and Weber, 2024).

The analysis consists of adjusting mathematical equations to data on leaf area (LA), total dry matter (TDM) and leaf dry matter (LDM), collected throughout the plants’ life cycle. The best equations (highest correlation coefficient and significant adjustment by F test) are chosen to estimate the physiological indicators of growth in instantaneous values: crop growth rate (CGR), relative growth rate (RGR), net assimilatory rate (NET), leaf area ratio (LAR), specific leaf area (SLA) and leaf area duration (LAD).

As an additional option in growth analysis, this program was developed, making some analyzes easier and faster for comparing different genotypes or for genotype and environment interaction studies. To carry out the growth analysis with the proposed program, data on leaf area, total and leaf dry matter and the respective collection dates or age of the plants must be available.

This program is an updated and improved version of the Anacres program created in 1990 in the Basic language (Portes and Castro, 1991). To access it, go to www..... (to be inserted). In developing the program, the C* language (C Sharp) from the Visual Studio platform was used.

## Material and methods

Calculation of instantaneous growth indicators using equations adjusted to the data.

In this work, the main equations that best fit plant growth data were used. The development of the work included: adjustment of the equations to the collected data, determination of correlation coefficients (r) and the F test (ANOVA). With the calculated F value available, the significance of the adjustment is found by searching for the F value in the table, usually as annexes in statistics books (Larson and Farber, 2010). The adjustment is significant when the calculated F value is greater than the tabulated F value. To find the tabulated F value, enter the regression degree of freedom (DF) value in the horizontal line of the table and the residual degree of Freedom in the column.

The analysis is based on fitting the equations to the data using the least squares method (Spiegel, 1985). Equations were adjusted to data on leaf area (FA), total dry matter (MST) and leaf dry matter (MSF), as a function of plant age (t) in days after emergence (DAE). The systems of linear equations obtained were solved by determinants, using the methods of Kramer, Sarrus and Leibniz (Landsberg, 1977; Dorn and McCracken, 1978; Spiegel, 1985; Ruggiero and Lopes, 1988; Pereira and Arruda 1987). The equation that best fits the analyzed data is the one with the highest correlation or determination coefficient and the fit, significant by the F test

Once the best equations for the LA, TDM and LDM data have been found, they will be used to determine the instantaneous growth indicators, applying derivative: crop growth rate (CGR = dTDM/dt; where d is the derivative of the equation TDM as a function of age t = DAE), relative growth rate (RGR= (dTDM/dt)/TDM), The relative growth rate at any instant is defined as the rate of increase in biomass per unit of biomass present: net assimilatory rate (NAR = dTDM/dt)/LA), specific leaf area (SLA = LA/LDM), leaf area ratio (LAR=LA/TDM), leaf area duration 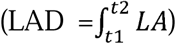 and fluxes of CO_2_ (Pn = Net Photosynthesis).

The equations adjusted to the leaf area (LA), total dry mass (TDM) and leaf dry mass (LDM) data, considered the dependent variables depending on the age of the plants (DAE), independent variable, were linear, sigmoid or logistic, quadratic, cubic, quadratic exponential and cubic exponential (Table 1).

**Table 1.**
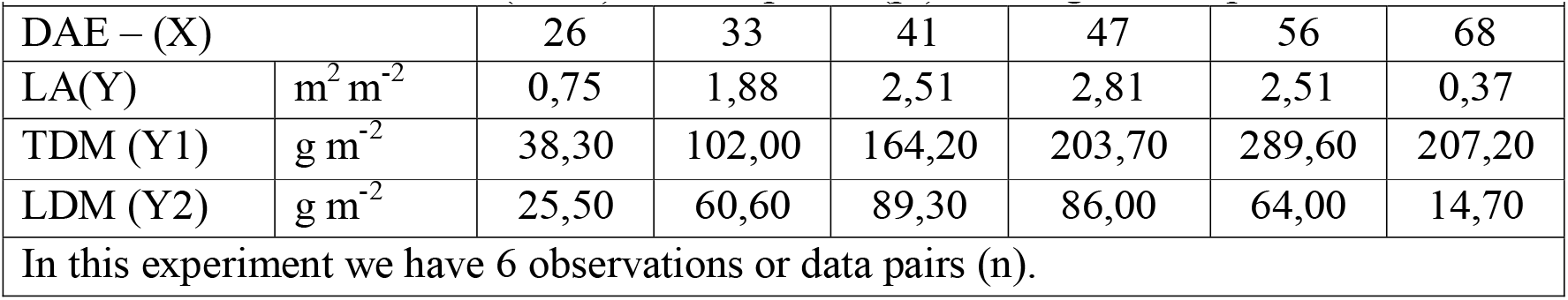
Values of leaf area (FLA), total dry matter (TDM) and leaf dry matter (LDM) at different collection dates (DAE) of bean plants (pl). Average of 4 repetitions.

Calculations of the average physiological growth indicators were also carried out and shown using the deduced classical equations found in plant physiology books and articles (Radford, 1967; KVET *et al*., 1971).

Calculation of average growth indicators using classical equations

The classic equations used to calculate average growth indicators are those found in the literature (Fisher, 1921; Radford, 1967; KVET *et al*., 1971):

Average crop growth rate 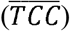. The dash over the acronym means average value.

The average dry matter production rate or average growth rate in a time interval T1 to T2 is given by the equation:

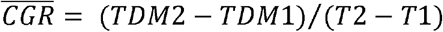

The mean Relative Crop Growth Rate 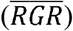 is defined as the rate of increase in biomass per unit of biomass present and is given by the equation (Fisher, 1921):

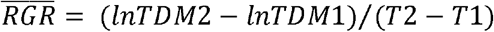

The net assimilation rate 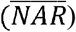, expresses the rate of increase in dry matter between two dates (T1, T2) in relation to the leaf area. The leaf area (LA) representing an estimate of the size of the assimilatory apparatus.

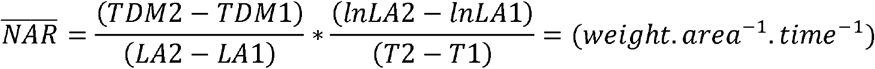

Leaf area duration is the sum of the leaf area of the plant throughout its cycle or part of it.

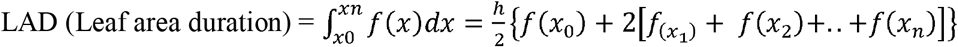

The integration was carried out using the trapezoidal summation method, with h being their height and f(x) being the result of the equation for each value of x (DAE).

Testing the program. To test the program, we analyzed the data presented in Table 1. The table contains data on leaf area (LA) in m^2^ per plant (m^2^ pl^-1^), total dry matter (TDM) and leaf dry matter (LDM) in grams per plant (g pl^-1^). These data were collected in an experiment conducted in a greenhouse at the Institute of Biological Sciences (ICB) at the Federal University of Goias (UFG). The experiment was conducted in pots with a capacity of 10 L of fertilized soil according to soil chemical analysis.

The first step in the program is to enter the data in Form 1. All are entered in the first form for each collection date.

Once the data has been entered and saved, move on to the next form. In this, equations will be adjusted to the LA, TDM and LDM data depending on the collection dates or sampling dates (DAE = T) (Table 2). Once the best equations have been found, we move on to the next form, which contains options for calculating physiological growth indicators (Table 3).

**Table 2.** Equations adjusted to LA, TDM and LDM data (y) depending on the collection dates - DAE (X) and their versions for adjustment.

|  |  |  |
| --- | --- | --- |
| Equation |  | Version to adjust the equation |
| Linear | $y = a_0 + a_1X$ | $y = a_0 + a_1X$ |
| Logistic | $y = C/(1+A*\exp(-a_1*X))$ | $\ln (C/y-1) = \ln A - BX$ Para $y > 0, y < C$ |
| Quadratic | $y = a_0 + a_1X + a_2X^2$ | $y = a_0 + a_1X + a_2X^2$ |
| Cubic | $y = a_0 + a_1X + a_2X^2 + a_3X^3$ | $y = a_0 + a_1X + a_2X^2 + a_3X^3$ |
| Quadratic Exponential | $y = c*\exp*(a_1X + a_2X^2)$ | $\ln y = \ln c + a_1X + a_2X^2$<br>faz-se $\ln y = Z$ e $\ln c = a_0$ |
| Cubic Exponential | $y = c*\exp*(a_1X + a_2X^2 + a_3X^3)$ | $\ln Y = \ln c + a_1X + a_2X^2 + a_3X^3$<br>faz-se $\ln y = Z$ e $\ln c = a_0$ |

**Table 3.**
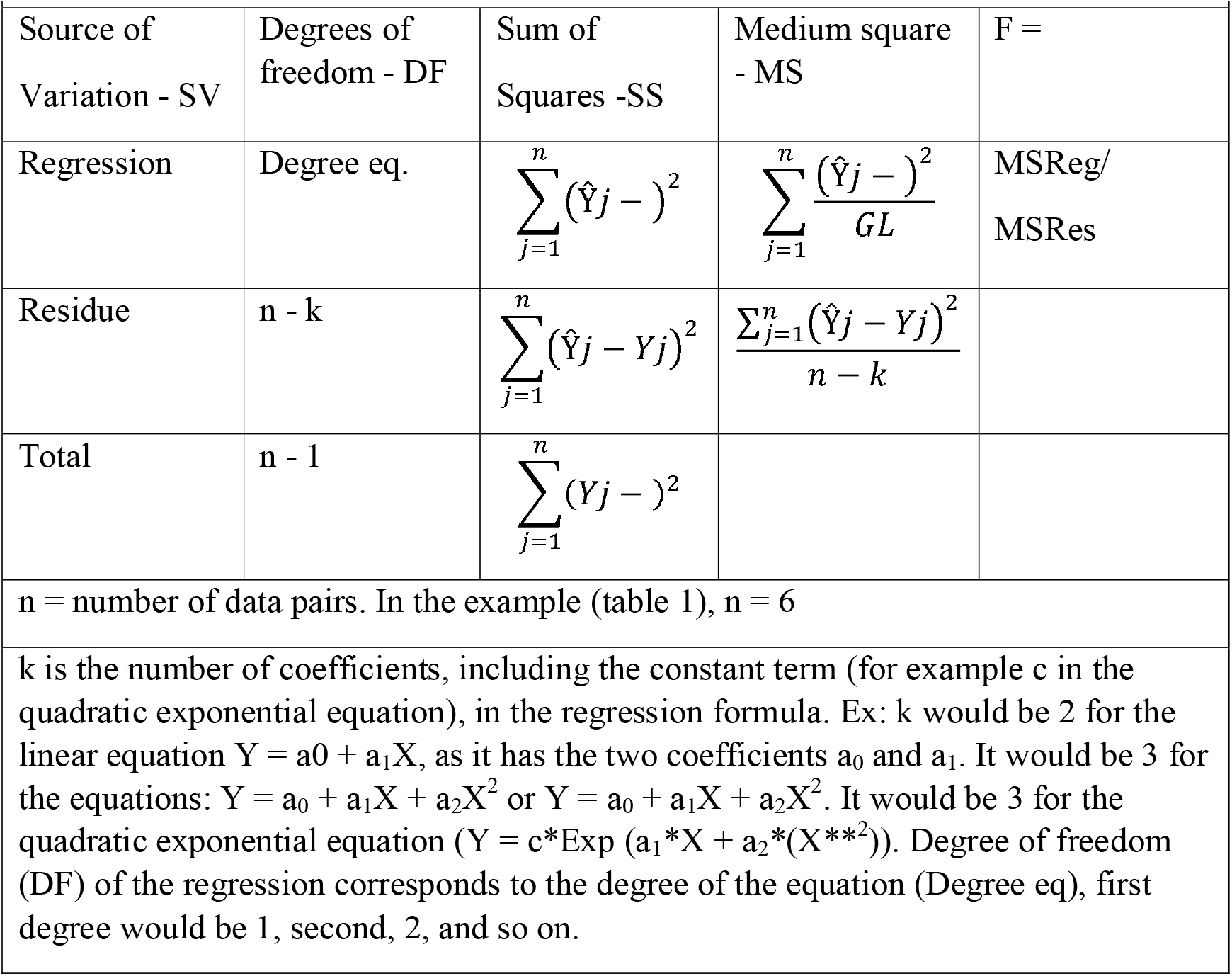
ANOVA for regressions.

| **Table 3.** ANOVA for regressions |  |  |  |  |
| --- | --- | --- | --- | --- |
| Source of Variation - SV | Degrees of freedom - DF | Sum of Squares -SS | Medium square - MS | F = |
| Regression | Degree eq. | $\sum_{j=1}^n (\hat{Y}_j - )^2$ | $\sum_{j=1}^n \frac{(\hat{Y}_j - )^2}{GL}$ | MSReg/<br>MSRes |
| Residue | n - k | $\sum_{j=1}^n (\hat{Y}_j - Y_j)^2$ | $\frac{\sum_{j=1}^n (\hat{Y}_j - Y_j)^2}{n - k}$ | |
| Total | n - 1 | $\sum_{j=1}^n (Y_j - )^2$ | | |
| n = number of data pairs. In the example (table 1), n = 6 |  |  |  |  |
| k is the number of coefficients, including the constant term (for example c in the quadratic exponential equation), in the regression formula. Ex: k would be 2 for the linear equation $Y = a_0 + a_1X$, as it has the two coefficients $a_0$ and $a_1$. It would be 3 for the equations: $Y = a_0 + a_1X + a_2X^2$ or $Y = a_0 + a_1X + a_2X^2$. It would be 3 for the quadratic exponential equation ($Y = c \cdot \text{Exp}(a_1 \cdot X + a_2 \cdot (X^{**2}))$). Degree of freedom (DF) of the regression corresponds to the degree of the equation (Degree eq), first degree would be 1, second, 2, and so on. | | | | |

The calculations of the coefficients or parameters of the linear, logistic, quadratic and quadratic exponential equations were carried out using determinants: The matrix was constructed and the Kramer rule (Spiegel, 1984; Pereira and Arruda, 1987) and the Sarrus rule were used. For the cubic and cubic exponential equations, the coefficients were calculated using the rule of Leibniz. (https://matrix.reshish.com/ptBr/cramer.php).

The correlation coefficient r is calculated by the square root of the relationship between the explained variation and the total variation.

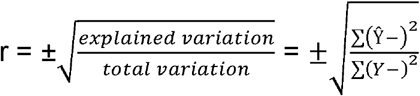

where Ŷ is the Y estimated or calculated by the adjusted equation, □ is the average value of the Y values observed in the laboratory or in the field.

This equation for r can be applied to both linear (such as linear regression) and non-linear (quadratic, cubic, quadratic exponential, cubic exponential) functions (Spiegel, 1984).

The analysis of variance was carried out according to Table 3.

Once the equations have been adjusted and the r and F values have been calculated, they are used to estimate the physiological indicators of plant growth: Leaf area ratio and Specific leaf area. The other growth indicators are estimated by deriving the equations to find the Instantaneous Crop Growth Rates (CGR), relative growth rates (RGR), Net Assimilation Rate (NET) and CO_2_ Flow Rate (Table 4 and 5).

**Table 4.**
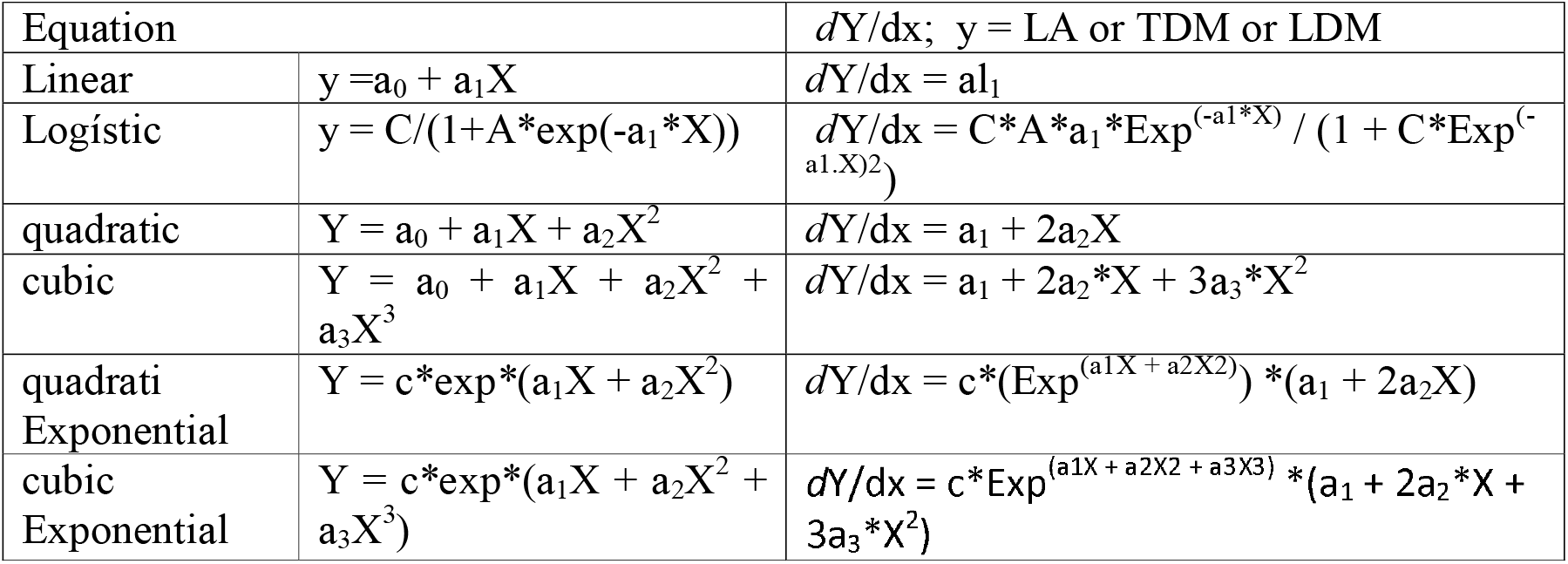
Equations adjusted to leaf area (LA), total dry mass (TDM) and leaf dry mass (LDM) data and their respective derivatives. X = collection dates in the Julian calendar (days after plant emergence).

**Table 5.** Plant growth indicators estimated from the equations adjusted to the data.

|  |  |  |  |  |  |
| --- | --- | --- | --- | --- | --- |
| CGR | RGR | NET | LAR | SLA | Pn |
| $dTDM/dt$ | CGR/<br>TDM | CGR/<br>LA | LA/TDM | LA/ADM | $P_n = NET * \left(\frac{C}{100}\right) * \left(\frac{44}{12}\right) * \left(\frac{1}{44}\right) * \left(\frac{1}{86400}\right) * (10^6)$ |
| $P_n = NET * 0,434027778$ para C = 45% | | | | | |

## Results and discussion

The equations that best fit the data on leaf area, total and leaf dry matter were quadratic, cubic, quadratic exponential and cubic exponential, based on their correlation coefficients and significance levels (Table 6).

**Table 6.** The best equations adjusted to the LA, TDM and ADM data, depending on the correlation coefficients and significance using the F test.

| Equation | Leaf area | r | F | p |
| --- | --- | --- | --- | --- |
| Quadratic | $y = -8,12327487 + (0,47108674 X) + (-0,00507602 X^2)$ | 0,99 | 129,3 | $\leq 0.01$ |
| Cubic | $y = -4,06317818 + (0,17819644 X) + (0,0015283 X^2) + (-0,00004689 X^3)$ | 0,99 | 242,9 | $\leq 0.01$ |
| Quadratic Exponential | $y = 0,00101187765110373 * (\text{Exp}(0,35598118 X) + (-0,00394095 X^2))$ | 0,98 | 48,2 | $\leq 0.01$ |
| Exponential Cubic | $y = 0,04564117 * \text{Exp}(+ (0,08115142 X) + (0,00225613 X^2) + (-0,00004399 X^3))$ | 0,99 | 35,9 | $\leq 0.05$ |
| Total Dry Mass |  |  |  |  |
| Quadratic | $y = -478,30540864 + (24,92636353 X) + (-0,21470086 X^2)$ | 0,96 | 16,9 | $\leq 0.05$ |
| Cubic | $y = 451,39233206 + (-42,14087082 X) + (1,29758451 X^2) + (-0,01073603 X^3)$ | 0,97 | 31,8 | $\leq 0.01$ |
| Quadratic Exponential | $y = 0,40535 * \text{Exp}(0,23062018 X) + (-0,00204173 X^2)$ | 0,99 | 96,3 | $\leq 0.01$ |
| Cubic Exponential | $y = 0,14546306 * \text{Exp}(+ (0,30454982 X) + (-0,00370875 X^2) + (0,00001183 X^3))$ | 0,95 | 17,8 | $\leq 0.01$ |
| Leaf Dry Mass |  |  |  |  |
| Quadratic | $y = -230,38631237 + (13,8216298 X) + (-0,15081952 X^2)$ | 0,99 | 71,8 | $\leq 0.01$ |
| Cubic | $y = -375,0570393 + (24,2579955 X) + (-0,38614699 X^2) + (0,00167064 X^3)$ | 0,99 | 79,5 | $\leq 0.01$ |
| Quadratic Exponential | $y = 0,07664 * \text{Exp}(0,31588983 X) + (-0,0035079 X^2)$ | 0,99 | 304,7 | $\leq 0.01$ |
| Cubic Exponential | $y = 0,0452461 * \text{Exp}(+ (0,35390774 X) + (-0,00436516 X^2) + (0,00000609 X^3))$ | 0,99 | 51,3 | $\leq 0.01$ |

The values observed and calculated using the equations adjusted to the data are below (Fig. 1,2,3):

**Fig. 1.**
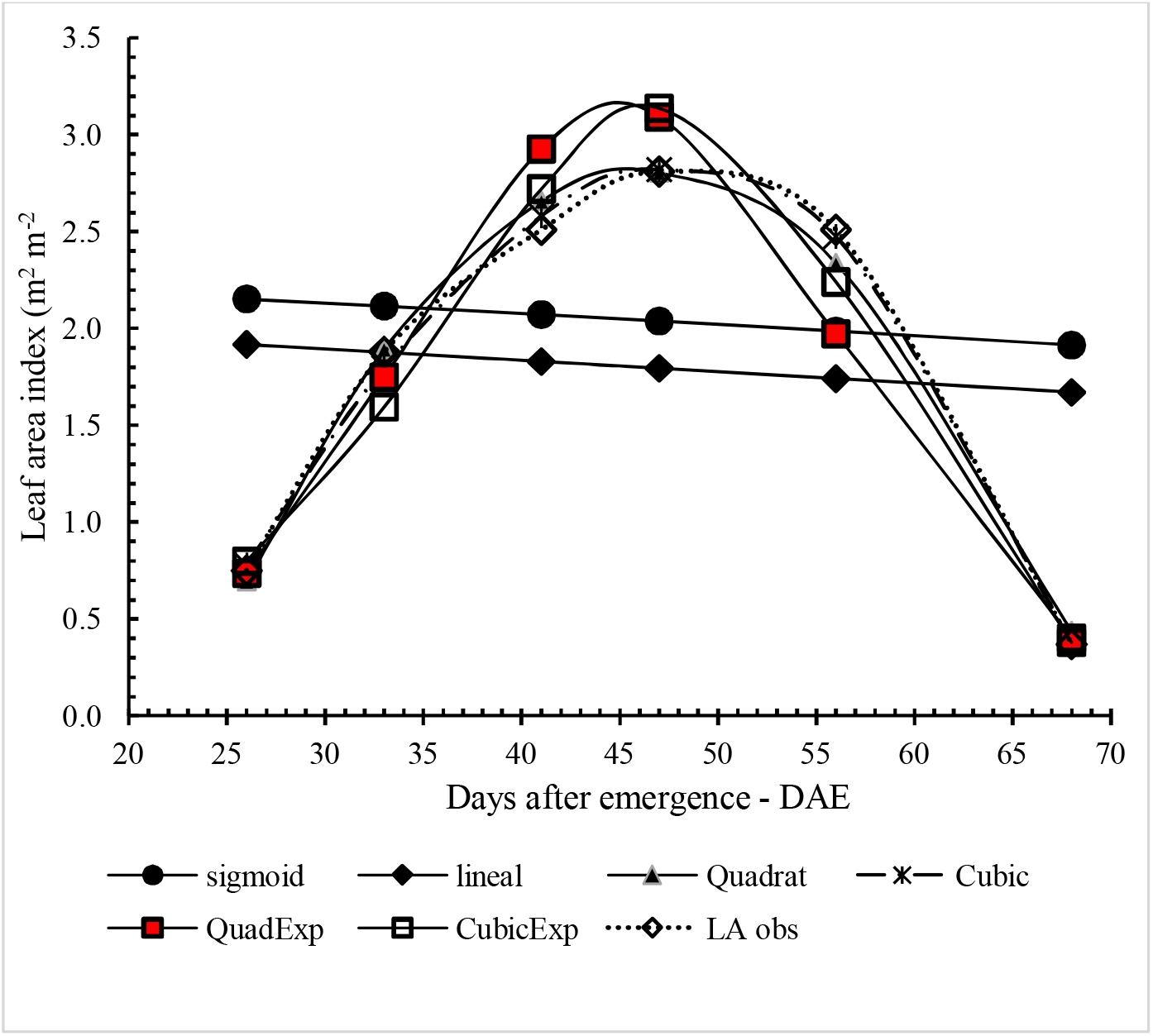
Curves showing the observed (LAobs) and calculated values of leaf area index using the equations fitted to the data.

**Fig. 2.**
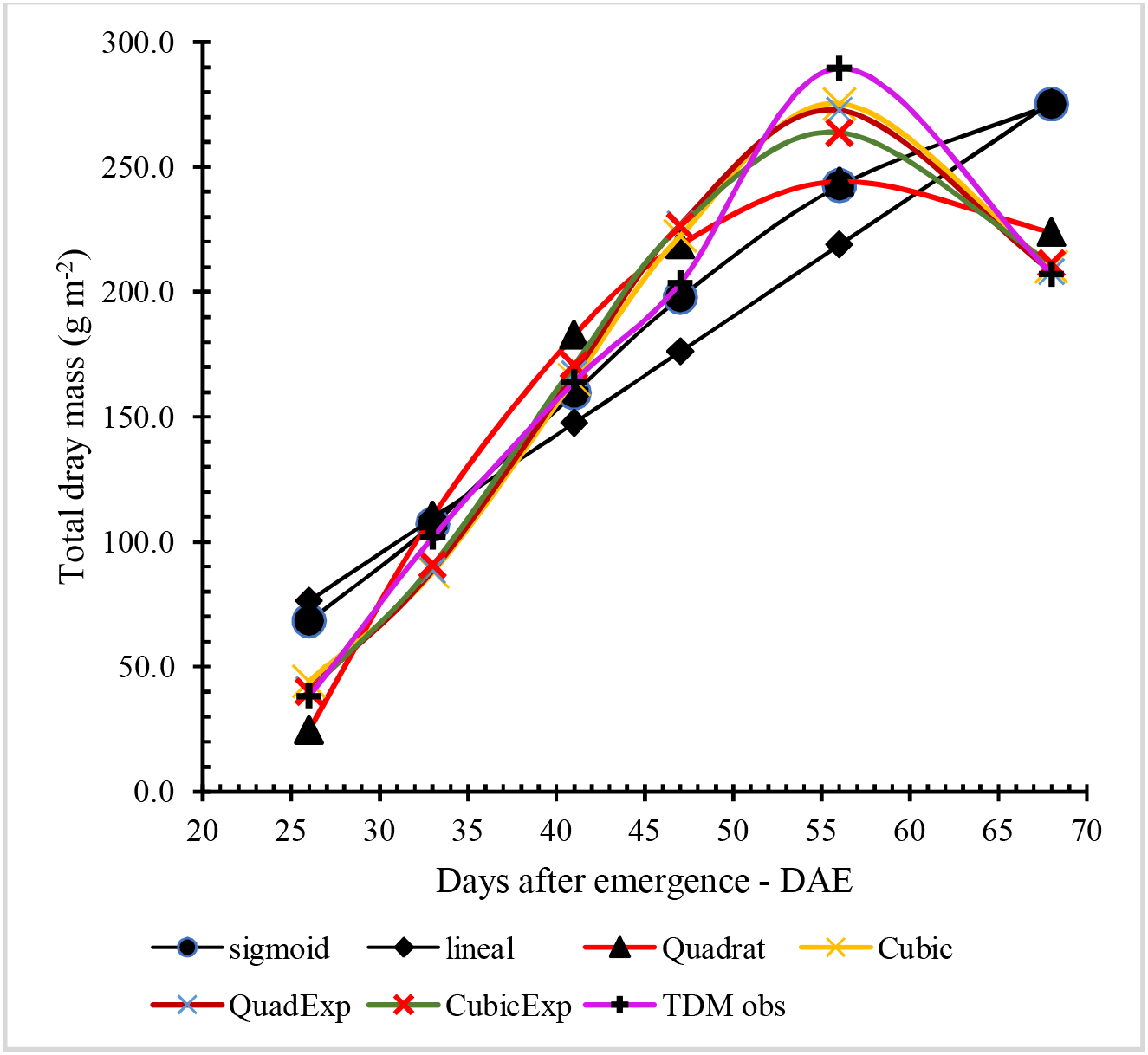
Curves showing the observed (TDMobs) and calculated values of total dry mass using the equations fitted to the data.

**Fig. 3.**
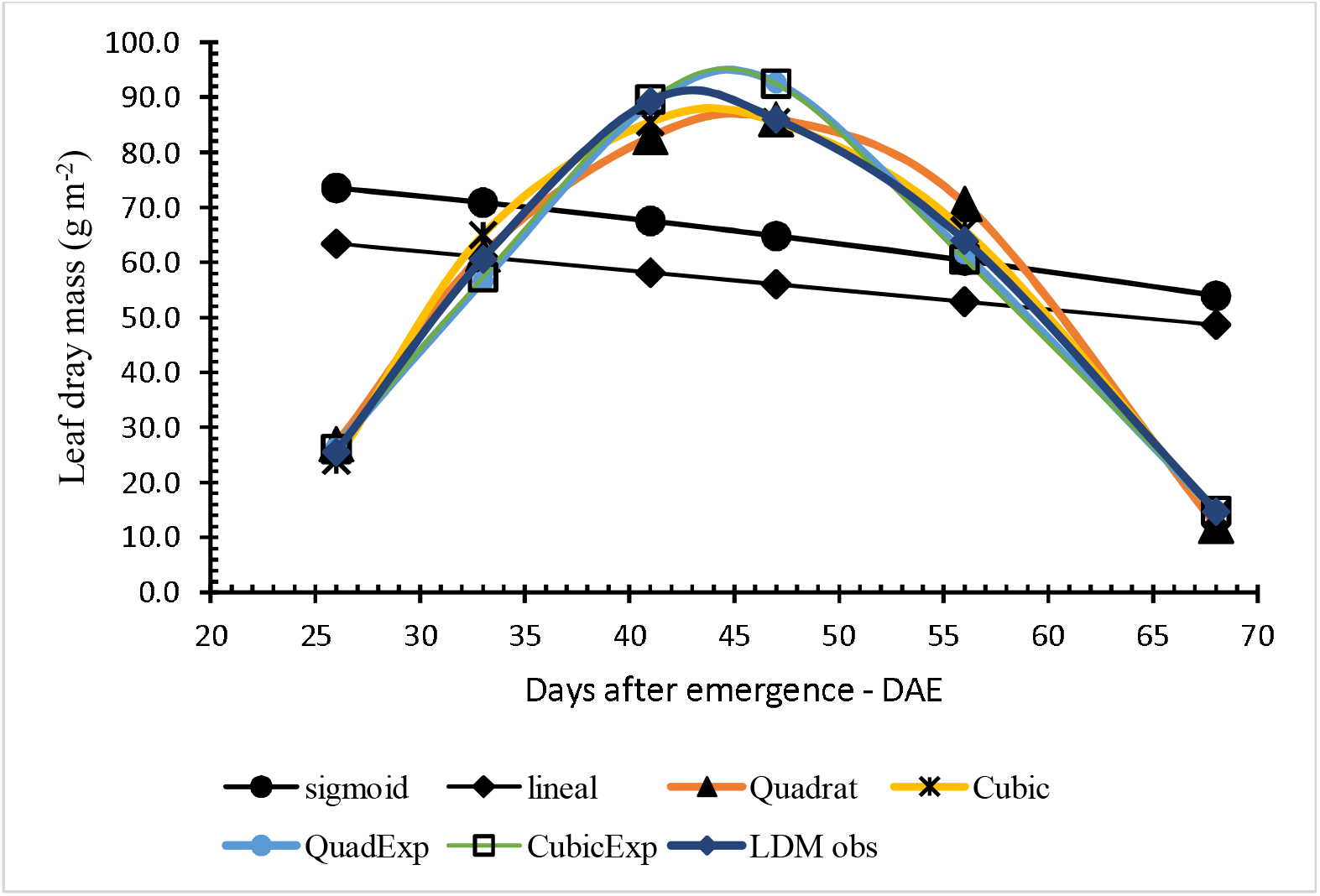
Curves showing the observed (LDM obs) and calculated values of leaf dry matter using the equations fitted to the data.

### Growth indicators

The growth indicators in instantaneous values (CGR, RGR, NET, LAR, SLA and Pn) were obtained using the equations presented in Table 7.

**Table 7.**
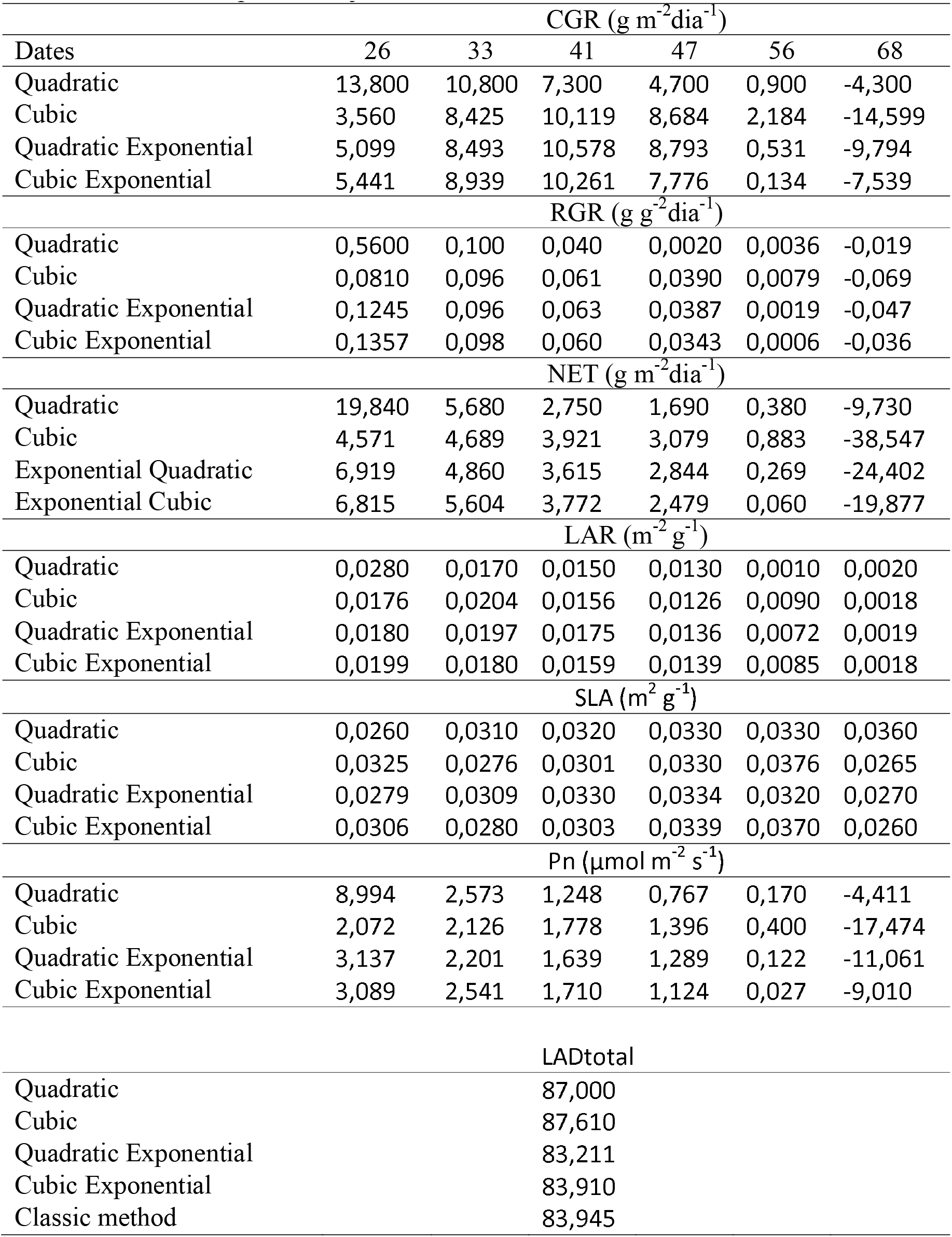
Instantaneous growth indicators CGR, RGR, NET, LAR, SLA, LAD and Pn estimated from the equations adjusted to the data.

Table 8 contains the values of the mean growth indicators 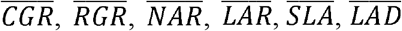 and Pn estimates from classical equations

**Table 8.**
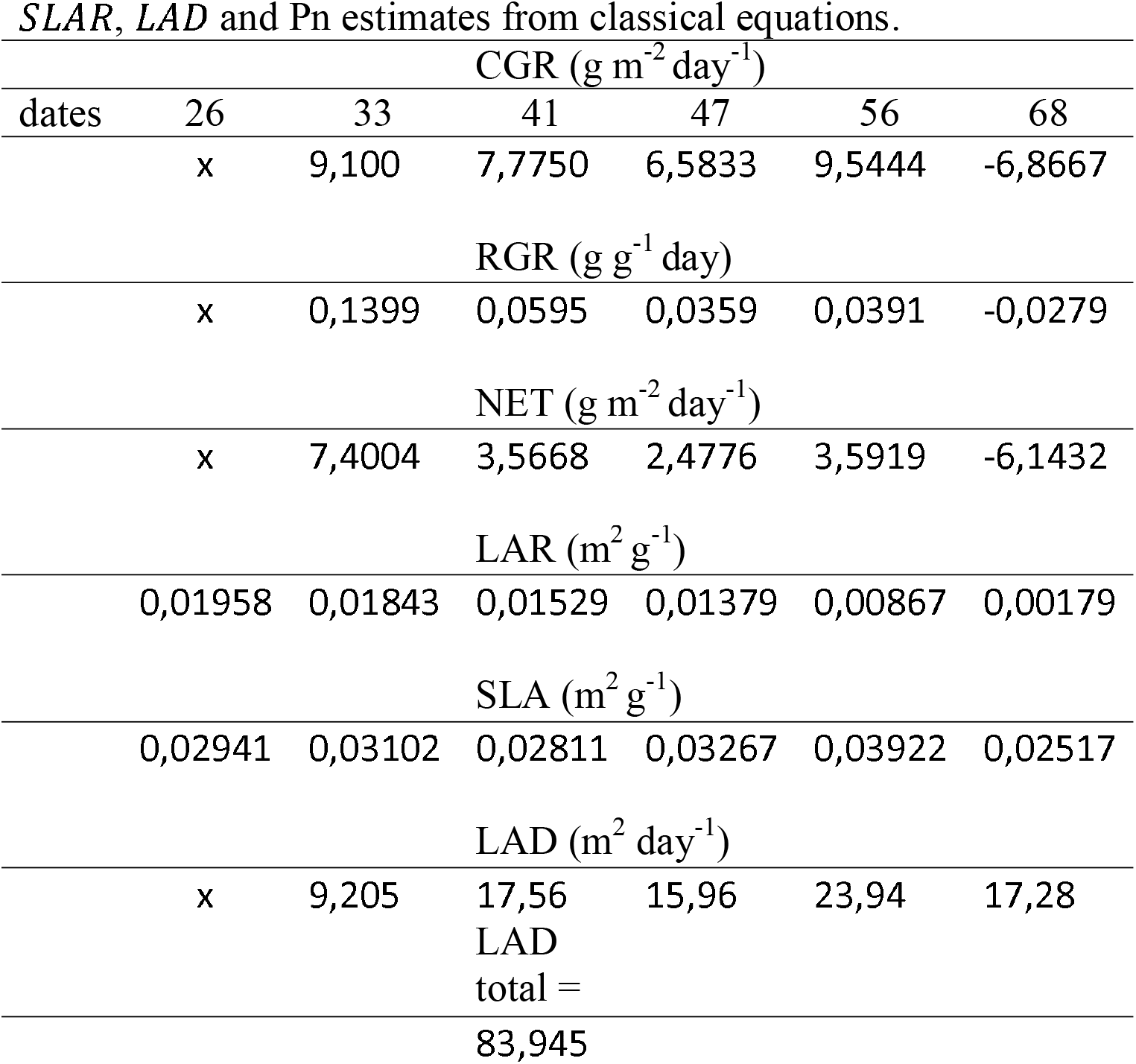
Average growth indicators 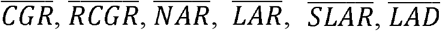 and Pn estimates from classical equations.

It is interesting to compare the results obtained for the growth indicators in instantaneous values and those estimated by classical equations. To facilitate visualization, the graphs are presented in Figure 4. The differences observed indicate that the values obtained from the equations adjusted to the LA, TDM and LDM data are more reliable due to the high correlation coefficients and significance found for the aforementioned equations.

**Fig. 4.**
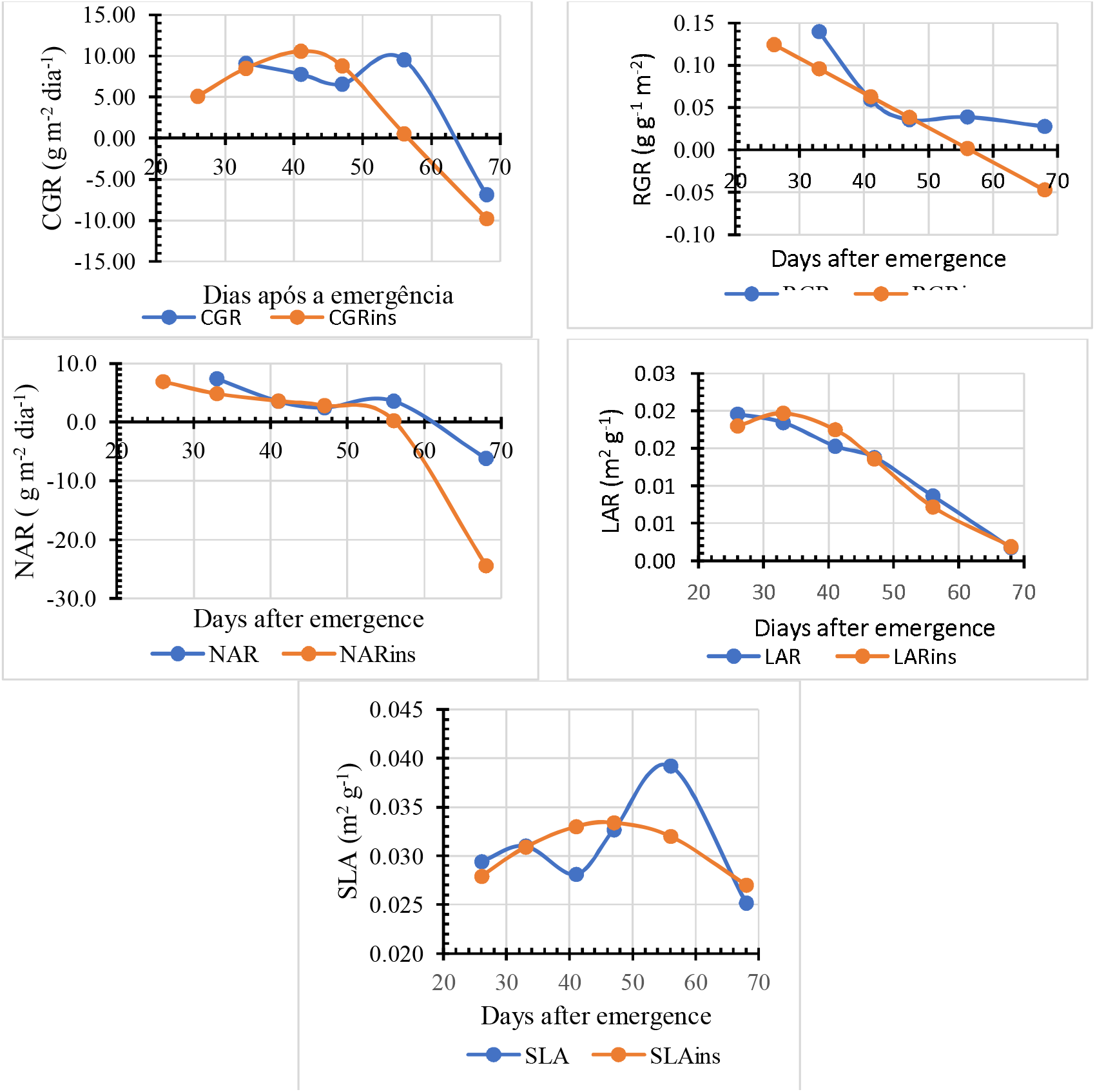
Comparisons between growth indicators calculated using classical formulas (mean values) and those estimated from equations adjusted to data from TDM, LA, LDM (instantaneous values). Crop Growth Rate (CGR), Relative Growth Rate (RGR), Net Assimilate Rate (NAR), Leaf Area Rate (LAR), Specific Leaf Area Rate (SLR). (ins = instantaneous values)

## Conclusions

Growth analysis is a classic technique, but it is still important due to the need to measure plant growth whenever they are subjected to some treatment, which can be biotic or abiotic. It is through photosynthesis that all organic matter is produced resulting in the development and growth of plants. Measuring photosynthesis is a technique widely used to estimate the productive efficiency of plants in carbon assimilation, but the measurements are limited to quantifying the flow of CO_2_ to a restricted area of the leaf and, when extrapolating to a larger area, large errors can occur. The growth analysis is carried out by collecting several plants from the experimental area and the calculated averages of the analysed variables normally present low errors or coefficients of variation. This work presents an auxiliary computer program that is very useful in growth analysis calculations. It was observed that equations such as the quadratic exponential and the cubic exponential adjusted to data on leaf area, total dry matter and leaf dry matter showed high correlation coefficients and significance using the F test. As a result, the estimated growth indicators are very real, therefore reliable. The program facilitates the researcher’s work due to the agility of processing, being able to work with several treatments simultaneously. The program has the option of calculating growth indicators using the classic equations presented in the literature, but it is clear that the errors are large, justifying the use of equations adjusted to the data to obtain them.

## Abbreviations

CGR: crop growth rate
RGR: relative growth rate
NAR: net assimilate rate
SLA: specific leaf area
LAR: leaf area rate
LAD: leaf area duration
Pn: CO2 flux

### Appendix

Procedures used to adjust equations to data. The description is quite detailed to facilitate the understanding of the reader.

#### 1. Linear Equation

Equation: y = a_0_ + a_1_X. The system of equations for the linear equation is as follows (Spiegel, 1984, https://matrix.reshish.com/ptBr/cramer.php):

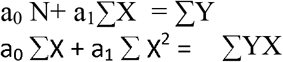

With a_0_ and a_1_ being the coefficients of the equation, N is the number of data pairs or number of collections, ∑X is the sum of the values of X, ∑Y is the sum of the values of Y e ∑YX is the sum of the products of Y times X.

From the system of equations, the matrix is obtained from which the coefficients a_0_ and a_1_ are found:

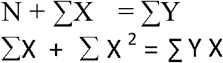

To find the coefficients a_0_ and a_1_, Kramer’s rule is applied (Spiegel, 1984).

(Det A).

Det A = N* ∑X^2^ - ∑X * ∑X; Det a0 = ∑Y * ∑X^2^ - ∑X * ∑YX; ao = Det a0/Det A;

Det a_1_ = N * ∑YX - ∑Y * ∑X; a1 = Det a1/Det A.

Once the values of the coefficients a_0_ and a_1_ have been found, just replace them in the equation.

#### 2. Logistic Equation

To fit the equation to the data it is necessary to find the values of C, A and a1

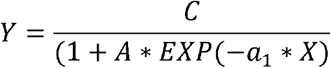

To make it easier, first transform the equation into a linear one, then the previous equation looks like this:

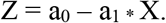

How to proceed?

Transferring the C to the left, it becomes:

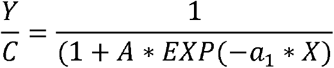

Inverting the numerator with the denominator to make it easier:

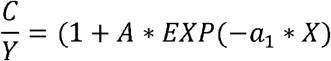

Now transferring 1 to the left of the equation:

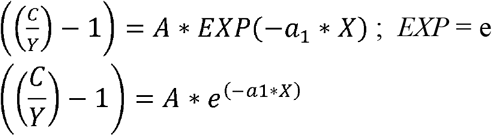

And applying logarithm (basic logarithm rules):

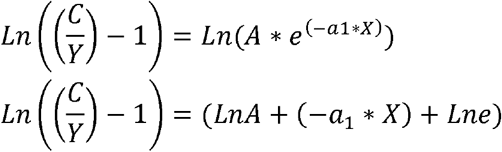

Lne = 0 logaritmo de e na base e = zero, então fica:

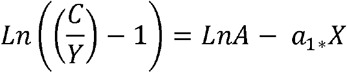

Doing: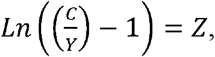

and Ln A = a_0_

The equation looks like the linear one (already seen):

Z = a_0_ – a_1_ * X. Once the values of a_0_ and a_1_ are found, it becomes possible to find the values of C and A from the logistic equation. The value of C cannot be equal to the value of maximum Y, because it would be C = maximum Y and C/ maximum Y = 1 and 1 – 1 = 0, and there is no logarithm of Zero. To make it easier, let’s make C = ymaximo * 102/100, this way, C will always be slightly higher than Ymaximum (102/100 is an arbitrary constant, used only to make C a value slightly higher than Ymaximum). A = e^ao^.

#### 3. Quadratic Equation

The quadratic equation is as follows: Y = a_0_ + a_1_X + a_2_X^2^. Adjusting this equation to the data (LA, TDM and LDM) means finding the values of the constants or parameters of the equation: a_0_, a_1_ and a_2_. How to proceed? First, the system of equations is set up (Spiegel, 1985).

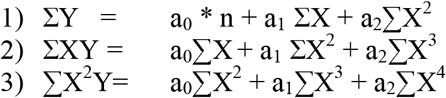

From this system of equations we obtain the matrix

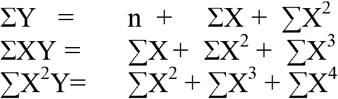

The coefficients a_0_, a_1_ and a_2_ of the equation were obtained using determinant and Kramer’s rule (Spiegel, 1985).

Calculation of the determinant A (DetA)

DetA = [(N * ∑x2 * ∑x4) + (∑x * ∑x3 * ∑x2) + (∑x2 * ∑x * ∑x3)] - [(∑x2 * ∑x2 * ∑x2) + (N * ∑x3 * ∑x3) +(∑x * ∑x * ∑x4)]

Calculation of the determinant a_0_ (Det a_0_) and the coefficient a_0_

Deta0 = [(∑y + ∑x2 * ∑x4) + (∑x * ∑x3 * ∑yx2) + (∑x2 * ∑xy * ∑x3)] - [(∑x2 * ∑x2 * ∑yx2) + (∑y * ∑x3 * ∑x3) +(∑x * ∑yx * ∑x4)]

a_0_ = deta_0_ / detA;

Calculation of the determinant a_1_ (Det a_1_) and the coefficient a_1_

Deta_1_ = [(N * ∑y * ∑x4) + (∑y * ∑x3 * ∑x2) +(∑x2 * ∑y * ∑yx2)] - (∑x2 * ∑xy * ∑x2) +(N * ∑x3 * ∑yx2) +(∑yx * ∑x * ∑x4)]

a_1_ = deta_1_ / detA;

Calculation of the determinant a_2_ (Det a_2_) and the coefficient a_2_ Deta_2_

Deta_2_ = [(N * ∑x2 * ∑yx2) + (∑x * ∑yx * ∑x2) +(∑y * ∑x * ∑y)] - [(∑x * ∑x2 * ∑x2) + (N * ∑yx * ∑x3) +(∑x * ∑x * ∑yx2)]

a_2_ = deta_2_ / detA;

#### 4. Quadratic Exponential equation

The formula of the equation is 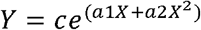. The adjustment consists of finding the values of the coefficients of equation c, a_1_ and a_2_. Applying logarithm: lneY = lne c + (_a1_ X + a_2_ X^2^) * ln_e_e. Solving: ln_e_Y = ln_e_c + (a_1_ X + a_2_ X^2^) x 1, since ln_e_e. In order to facilitate the adjustment, making lnY = Z and ln_e_c = a_0_, thus, the equation becomes as follow: Z = ao + a_1_ + a_2_ X^2^, solved in the same way as the quadratic. Finally, calculating c = e^a0^, simply substitute the coefficients in the original equation.

#### 5. Cubic equation

The cubic equation is as follows: Y = a_0_ + a_1_X + a_2_X^2^ + a_3_X^3^. The coefficients a_0_, a_1_, a_2_ and a_3_ of the equation will be obtained using determinant and Kramer’s rule and Sarrus’ rule (Spiegel, 1985, https://matrix.reshish.com/ptBr/cramer.php). The adjustment consists of finding the values of the coefficients of the equation a_0_, a_1_ and a_2_ and a_3_.

Calculate the determinant A (for simplicity the symbol ∑ was eliminated, so X2 is ∑X2, X4 is ∑X4 and so on).

detA = (n * X2 * X4 * X6) - (n * X2 * X5 * X5) - (n * X3 * X3 * X6) + (n * X3 * X5 * X4) + (n * X4 * X3 * X5) - (n * X4 * X4 * X4) - (X * X * X4 * X6) + (X * X * X5 * X5) + (X * X3 * X2 * X6) - (X * X3 * X5 * X3) - (X * X4 * X2 * X5) + (X * X4 * X4 * X3) + (X2 * X * X3 * X6) - (X2 * X * X5 * X4) - (X2 * X2 * X2 * X6) + (X2 * X2 * X5 * X3) + (X2 * X4 * X2 * X4) - (X2 * X4 * X3 * X3) - (X3 * X * X3 * X5) + (X3 * X * X4 * X4) + (X3 * X2 * X2 * X5) - (X3 * X2 * X4 * X3) - (X3 * X3 * X2 * X4) + (X3 * X3 * X3 * X3).

Calculate the determinant a_0_ and the coefficient a_0_

Deta_0_ = (Y * X2 * X4 * X6) - (Y * X2 * X5 * X5) - (Y * X3 * X3 * X6) + (Y * X3 * X5 * X4) + (Y * X4 * X3 * X5) - (Y * X4 * X4 * X4) - (YX * X * X4 * X6) + (YX * X * X5 * X5) + (X * X3 * YX2 * X6) - (X * X3 * X5 * YX3) - (X * X4 * YX2 * X5) + (X * X4 * X4 * YX3) + (X2 * YX * X3 * X6) - (X2 * YX * X5 * X4) - (X2 * X2 * YX2 * X6) + (X2 * X2 * X5 * YX3) + (X2 * X4 * YX2 * X4) - (X2 * X4 * X3 * YX3) - (X3 * YX * X3 * X5) + (X3 * YX * X4 * X4) + (X3 * X2 * YX2 * X5) - (X3 * X2 * X4 * YX3) - (X3 * X3 * YX2 * X4) + (X3 * X3 * X3 * YX3).

a_0_ = Deta_0_ / detA.

Calculate the determinant a_1_ and the coefficient a_1_

Deta_1_ = (n * YX * X4 * X6) - (n * YX * X5 * X5) - (n * X3 * YX2 * X6) + (n * X3 * X5 * YX3) + (n * X4 * YX2 * X5) - (n * X4 * X4 * YX3) - (Y * X * X4 * X6) + (Y * X * X5 * X5) + (Y * X3 * X2 * X6) - (Y * X3 * X5 * X3) - (Y * X4 * X2 * X5) + (Y * X4 * X4 * X3) + (X2 * X * YX2 * X6) - (X2 * X * X5 * YX3) - (X2 * YX * X2 * X6) + (X2 * YX * X5 * X3) + (X2 * X4 * X2 * YX3) - (X2 * X4 * YX2 * X3) - (X3 * X * YX2 * X5) + (X3 * X * X4 * YX3) + (X3 * YX * X2 * X5) - (X3 * YX * X4 * X3) - (X3 * X3 * X2 * YX3) + (X3 * X3 * YX2 * X3).

a_1_ = Deta_1_ / detA.

Calculate the determinant a_2_ (Deta2) and the coefficient a_2_

Deta_2_ = (n * X2 * YX2 * X6) - (n * X2 * X5 * YX3) - (n * YX * X3 * X6) + (n * YX * X5 * X4) + (n * X4 * X3 * YX3) - (n * X4 * YX2 * X4) - (X * X * YX2 * X6) + (X * X * X5 * YX3) + (X * YX * X2 * X6) - (X * YX * X5 * X3) - (X * X4 * X2 * YX3) + (X * X4 * YX2 * X3) + (Y * X * X3 * X6) - (Y * X * X5 * X4) - (Y * X2 * X2 * X6) + (Y * X2 * X5 * X3) + (Y * X4 * X2 * X4) - (Y * X4 * X3 * X3) - (X3 * X * X3 * YX3) + (X3 * X * YX2 * X4) + (X3 * X2 * X2 * YX3) - (X3 * X2 * YX2 * X3) - (X3 * YX * X2 * X4) + (X3 * YX * X3 * X3).

a_2_ = Deta_2_ / detA.

Calculate the determinant a_3_ (Deta_3_) and the coefficient a_3_

Detaa_3_ = (n * X2 * X4 * YX3) - (n * X2 * YX2 * X5) - (n * X3 * X3 * YX3) + (n * X3 * YX2 * X4) + (n * YX * X3 * X5) - (n * YX * X4 * X4) - (X * X * X4 * YX3) + (X * X * YX2 * X5) + (X * X3 * X2 * YX3) - (X * X3 * YX2 * X3) - (X * YX * X2 * X5) + (X * YX * X4 * X3) + (X2 * X * X3 * YX3) - (X2 * X * YX2 * X4) - (X2 * X2 * X2 * YX3) + (X2 * X2 * YX2 * X3) + (X2 * YX * X2 * X4) - (X2 * YX * X3 * X3) - (Y * X * X3 * X5) + (Y * X * X4 * X4) + (Y * X2 * X2 * X5) - (Y * X2 * X4 * X3) - (Y * X3 * X2 * X4) + (Y * X3 * X3 * X3).

a_3_ = Deta_3_ / detA.

#### 6. Cubic exponential equation

The formula of the equation is 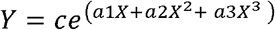. Applying logarithm, the equation becomes: lnY = ln c + (a_1_ X + a_2_ X^2^ + a_3_X^3^). Making lnY = Z and lnc = a_0_ the equation is as follows: Z = ao + a_1_ + a_2_ X^2^ + a_3_ X^3^, solved in the same way as the cubic one.

Finally calculating c = e^a0^, simply substitute the coefficients in the original equation.

